# Antibiotics and food in the American press

**DOI:** 10.1101/634337

**Authors:** Antoine Bridier-Nahmias, Estera Badau, Pi Nyvall-collen, Antoine Andremont, Jocelyne Arquembourg

## Abstract

The emergence of antimicrobial resistant infections from food is well documented in the scientific literature but, in this kind of matter, the public opinion is an important policy driver and is vastly forged by traditional media. Here, we propose a text mining study through about 500 articles from two reference daily U.S. newspapers to assess the media coverage of this issue. Our results indicate that, since the middle of the 80s, the two journals considered here adopted a very different narrative around the issue, echoing civil society concerns in one case and the official discourse in the other.

## Introduction

### Resistance to antibiotics is a growing concern

Microbial resistance to antibiotics (AMR) is a growing global public health concern. The grimmest projection even estimates the death toll due to this phenomenon to figures as high as 10 million per year by 2050 (***O’Neil Jim, 2016***), thus exceeding cancer-related deaths. The increasing public awareness on the subject has been pointed out by the World Health Organisation’s (WHO) 2014 report and its international action plan launched in 2015 in collaboration with the United Nation Food and Agriculture Organization (FAO) and the World Organisation for Animal Health (OIE) (***WHO, 2014, 2015***). The sanitarian actors alarm has also been followed by an international political commitment with the 41st G7 Summit of 2015, the 2016 “71st General Assembly of UN” and the 12th G20 Summit of 2017. Besides the need for international policies and standards on antibiotics use, these events show the political recognition of the links between the widespread use of antibiotics in human medicine and livestock, and the resistant pathogens transmission through the food chain (***Van Boeckel et al., 2015***). In this piece, we undertook a text mining study of the American press coverage of the link between antibiotic resistance and food. By comparing the New-York Times (NYT) and the Washington Post (WP) we were able to uncover a very different treatment of the subject that could have a strong influence on the public perception of the links between antibiotic resistance and food.

### Aim of the study

We decided to put the focus on the national American press coverage of antibiotic resistance impact on food through two reference newspaper with clear and distinct editorial lines: the New-York Times and the Washington Post. We implemented a text mining approach focusing on the scientific acknowledgment and the media coverage of the link between the antibiotic resistance and the transmission of resistant pathogens through the food chain. The phenomenon has been known and discussed since the middle of the 80s (***Holmberg et al., 1984***) and sparked debates in the media ever since. These debates revolve around exchanges between the scientific community, the political actors, the NGOs and the food-animal producers regarding the impact of antibiotic use in livestock on the loss of antibiotics’ efficacy and the need for regulation. Within the field of communication and media studies the subject is also gaining interest. A recent study by Carol Morris *et al.* analyses how in the UK the different actors in the debate on antibiotic use in agriculture defend their positions in the media (***Morris et al., 2016***). They worked on a corpus of 91 articles published by 4 national newspapers and one agricultural journal between 1998 and 2014. Another study of note by Bohlin and Host analyses the risk communication about antibiotic resistance in the Swedish daily press between 2008 and 2011 (***Bohlin and Höst, 2014***).

Our effort represents a first analysis of the American national press coverage of the link between antibiotic resistance and food through a systematic method that allows covering a broad period (1980-2016) and more than 500 articles, giving us access to social representations evolution regarding the subject and it’s media framing.

## Results

### Gradually rising coverage

Quantitatively, The New York Times number of publications surpasses The Washington Post (328 publications to 188 for 18,764 distinct words to 13,253 respectively used in the articles) and the researched subject gains a larger press coverage in both journals from 2010 onward (Fig. 1-A). As shown in figure 1-A, the similarity between the two newspapers regards the fact that the researched subject press coverage increases from 1980 to 2016, with peaks at some specific key moments. The peaks match different events related to the subject (Fig. 1-B). The latest peak (starting in 2010) is the most important in length and size. It corresponds to a multiple event sequence that could be divided as follows: the Food and Drug Administration (FDA) publishes a first draft regarding the “judicious use of antibiotics”, the announcement by the FDA of a voluntary plan to phase out antibiotics used as growth promoters in food-producing animals (2013), a nationwide episode of chicken contaminated with a drug-resistant salmonella (2013-2014), the release of a WHO report qualifying the antibiotic resistance a “global threat for human health” (2014) and the launch of the National Antibiotic Resistance Action Plan by the Obama Administration (2015) (Fig. 1-B). Clearly, the NYT’s coverage proportion of the researched subject has been higher than the WP’s since the first event.

**Figure 1.**
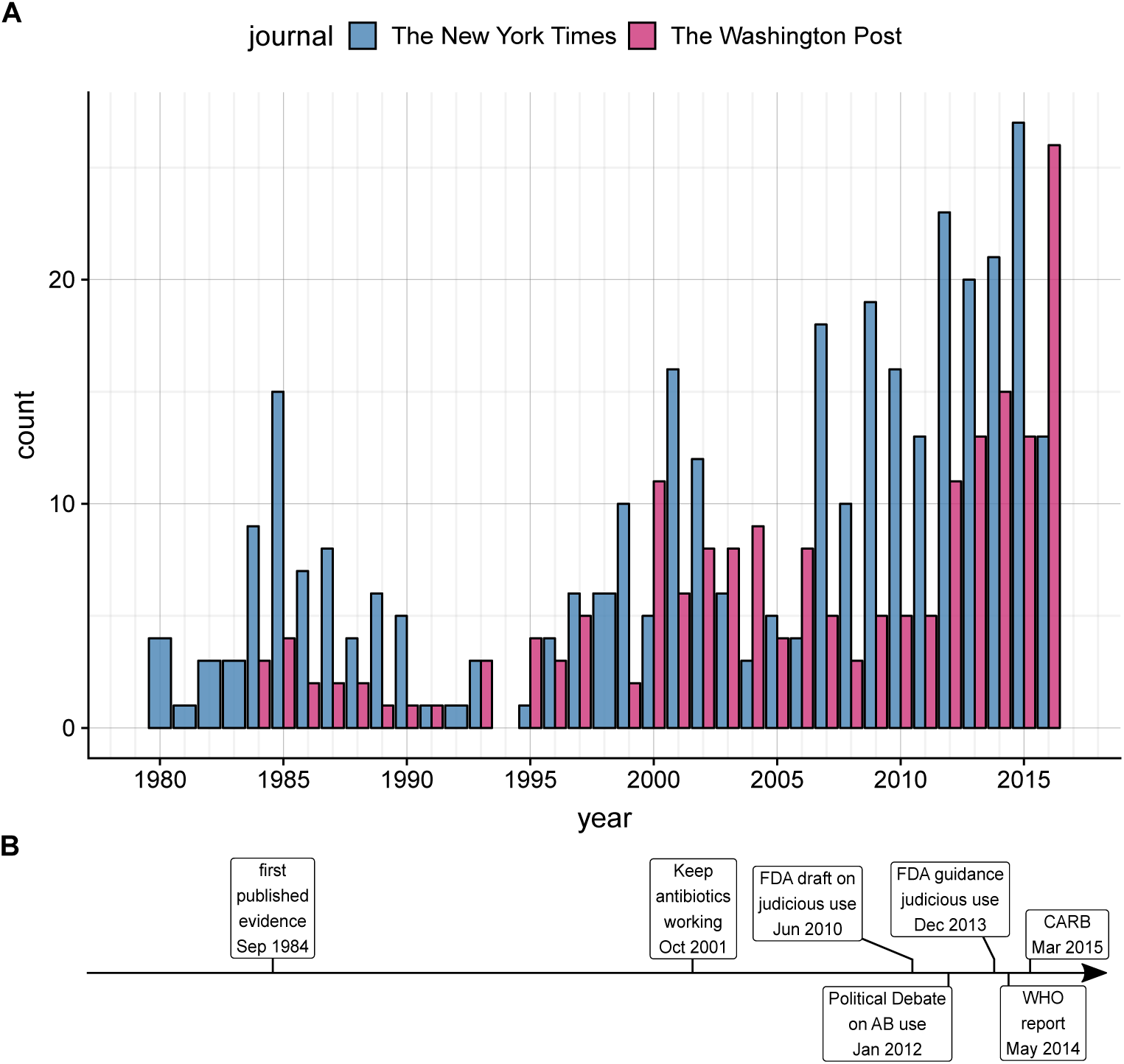
Evolution of the number of articles published from 1980 to 2016. The New York Times is represented in blue and the Washington Post in red. A: barplot with years on the x axis and the number of publication on the y axis. B: timeline focusing on a few key events. “first published evidence” refers to the publication of ***Holmberg et al.*** (***1984***), “Keep antibiotics working” is the creation of a coalition of various advocacy groups against the overuse of antibiotics in farming, “FDA draft on judicious use” issuance of the first draft guidance for the reduction of antibiotics usage, “Political debate on AB use” debate on the ban of some classes of antibiotics by the FDA ending with the ban of cephalosporins in animal feed, “FDA guidance judicious use” release by the FDA of the final version of their guidance for industry, “WHO report” publication of the global report on surveillance of antimicrobial resistance, “CARB”: president Obama issues the executive order CARB (Combating Antibiotic-Resistant Bacteria).

### Word usage across time

From the total of number of distinct words used in the whole corpus (22,880), the words with the highest number of occurrences and most used in the total number of the articles are, as expected from the search terms we used to gather the corpus, *antibiotics* (4004 occurrences) and *food* (3677), followed by *animals* (2523), *drug* (1892), *resistant* (1838), *farm* (1814), *organic* (1581), *meat* (1375), *health* (1354) and *chicken* (1237). Stop words (*i.e.* omnipresent and uninformative words) such as *the* or *and* were filtered out before counting using dictionary methods and manual curation.

The evolution of the most used terms by journal shows various patterns (Fig. 2). The use of the term *antibiotics* for example shows very similar trends between the two journals whereas the use of *farm* in the NYT shows multiple peaks but remains lower and steady in the WP. Likewise, Another term showing these differences between the two journals is agriculture. While in the WP publications its use was constant albeit pretty low (between 0 and 20 occurrences until 2014), the NYT publications use the term more frequently with several peaks between 20 and 60 occurrences (see Supp. data). A very different behavior is exhibited by the term industry. It shows a higher use by the NYT overall with some peaks appearing shortly after the events described in Fig. 1-B (1984: ‘First evidence’ & 2001: ‘Keep antibiotics working’). This is in constrast with the usage peaks visible in the WP in 1996, 1998, 2000 and 2004 that happen further away from events of our timeline. These discrepancies underline a very different usage of the term industry by the two journals.

**Figure 2.**
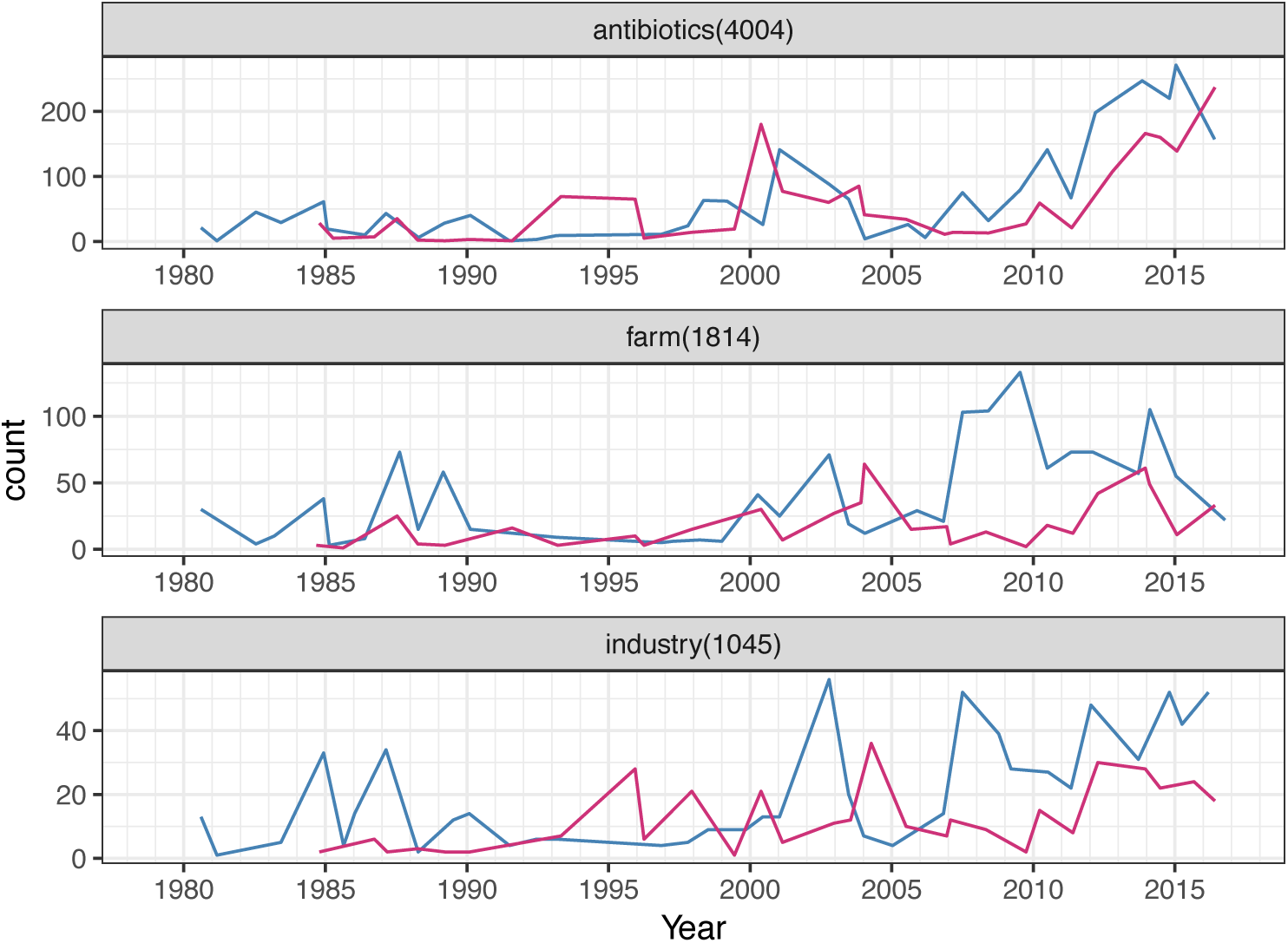
Yearly evolution of the use of selected terms. The year are represented on the x axis and the count on the y axis. The three examples chosen represent typical cases found in the corpus.

Patterns described above can be observed for many other terms and we can infer from these results that the two journals made different coverage choices regarding the events tied to antibiotic resistance and food supply. The NYT seems to put a stronger emphasis on agriculture, production and the food industry in general.

### *Antibiotic resistance* discursive context

The third part of our investigation regards the numbers of times a word was used within a sentence containing the expression *antibiotic resistance* (or one of its close variants). The results show that among the 299 sentences containing the expression, the nine most prominent terms were: *human* (72), *drugs* (68), *bacteria* (67), *animal* (64), *health* (59), *disease* (48), *food* (43), *public* (30) and *infections* (29). Interestingly, when we look at these results by journal, the NYT publications shows a higher occurrence for the following terms (among others): *food* (33 times in the NYT – 10 times in the WP), *salmonella* (10 – 5), *livestock* (12 – 5), *consumers* (5 – 2), *contaminated* (5 – 1), *outbreak* (4 – 1) and finally *poisoning* (4 – 0). Amidst the rare terms that are more frequently seen close to *antibiotic resistance* in the WP publications, we can note *FDA* (8 – 16), *agency* (4 – 9) and *officials* (2 – 10). These figures reveal a higher media coverage (beyond the sheer number of articles) of the direct link between the antibiotic resistance and food in the New York Times pages. They also hint at a higher coverage of the official discourse on the subject in the Washington Post.

### Discriminant terms between journals

After having looked thoroughly at the word counts, we decided to implement a machine learning strategy to highlight the differences between the NYT and the WP. We started by computing the term frequency - inverse document frequency (TF-IDF) of every word in each article (***Spärck Jones, 2004***). The TF-IDF can be considered here as a weight for each word characterizing an article inside the whole corpus.

Using these TF-IDF values as covariates, we fitted a cross-validated generalized linear model with an elastic-net penalization (***Friedman et al., 2010***) in order to identify the best subset of terms that allowed the classification of an article as being from the NYT or the WP. The resulting model (with an area under the ROC curve close to 0.83) retained seven variables (Fig. 3). The strongest predictors for the WP are the terms *cdc* and *rep* and to a lesser extent *launched*. The term rep refers to the mention of a member of the House of Representatives and *cdc* is the Center for Disease Control. On the NYT side, the predictive terms are *dr, editor, mr* and *ms*. Among them, editor comes mostly from the expression *letter to the editor*, indeed, when we analyzed the section in which the articles were published, we noted three times more letters in the NYT compared to the WP. What is clearly visible here is that the predictors associated with the WP are strongly related to the U.S. federal government whereas the NYT predictors are all directly linked to members of the civil society.

**Figure 3.**
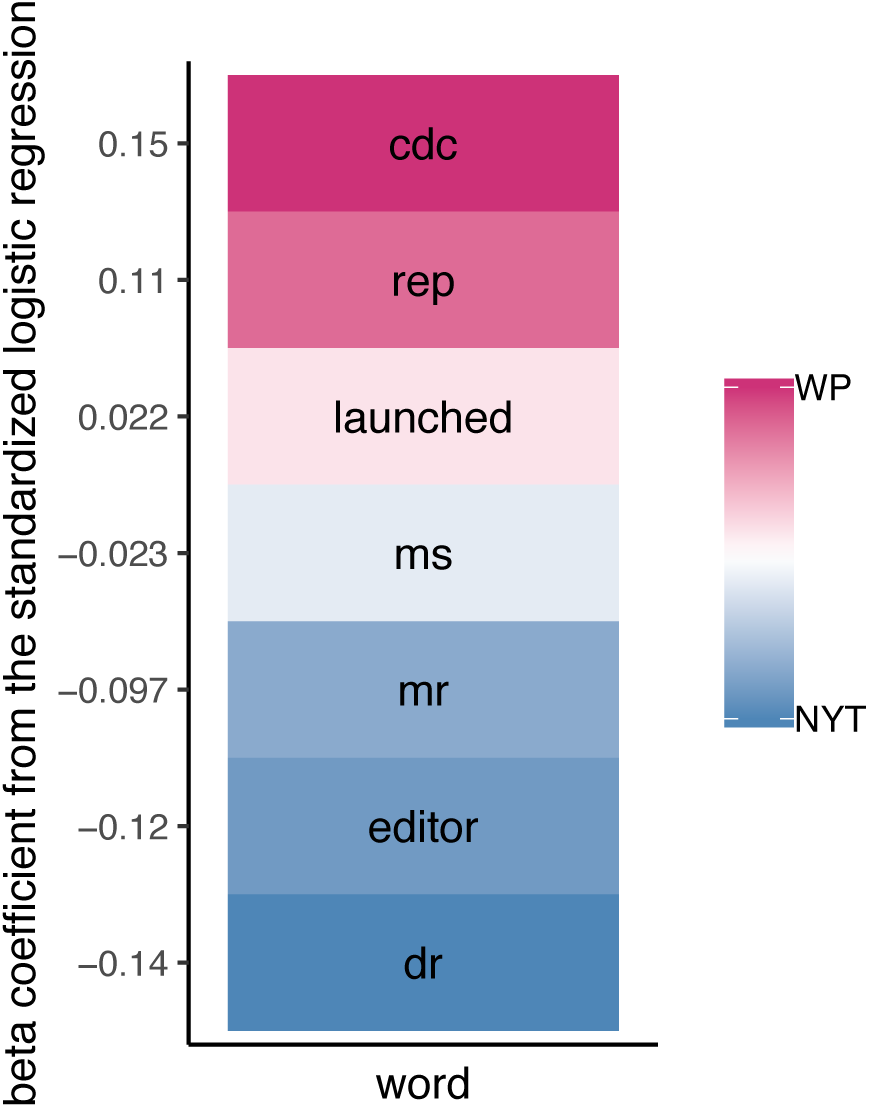
Words predicting the source of an article. The heatmap shows the beta coefficient of each word in the regression model. Positive values in red are associated with the WP, whereas negative values in blue are with the NYT. The numeric value represent the strength of the association.

## Conclusion

In this study we aimed at getting an overview of the collective representation of the problem of AMR and food in the US during the last three decades. In order to do so, we chose to analyze national daily press through two reference journals with different editorial lines. Through an analytical approach combining text-mining and frame analysis, we were able to uncover key differences in the media coverage of the issue. The most striking result was a clear difference between the discourse each journal chose to emphasize. On one hand, the NYT opens its pages to the civil society through quotes, citations and the publication of letters to the editor. On the other hand, the WP chooses to stand on the side of the institutional narrative of the debate, finding its sources in federal agencies such as the FDA, the CDC and the U.S Department of Agriculture. Aside from these general trends, we uncovered differences in term use through time and characterized three main patterns. The first group is constituted by very generic such as antibiotics that are used in a similar fashion in both journals. A second group of terms are massively used by one journal only and can be considered as a signature of their editorial line. Finally, the third group is constituted by terms seen in both journals but used asynchronously. The analysis of the last two groups described above reinforce our main finding pointing out the different media coverage of the subject. Finally, focusing on the sentences containing the expression *antibiotic resistance* allowed us to define the discursive context of the formula. A more in depth-look at these contexts by journal revealed differences concordant with our other results.

We restricted our study to written press to cover the whole period of interest starting in the 80s. In the future though, the methodology we defined could very easily scale-up and be applied to huge amounts of data scrapped from the internet. Moreover, access to textual resources from the social networks could provide a direct representation of the public awareness and participation in debate about policy involving social issue.

## Methods and Materials

Briefly, the articles were downloaded from the factiva database using multiple queries. The pdf were loaded into R (***R Core Team, 2019***) and parsed and transformed using r-base functions, tidyverse tools (***Wickham, 2017***) and the tidytext package (***Silge and Robinson, 2016***).

The complete analysis with embedded code in the form of an interactive html R notebook is available as supplementary data and online at the following address:https://a-bn.github.io/AntibioFoodTM/

## Acknowledgments

The authors would like to thank Alaksh Choudhury, Josselin Noirel and Marc Michel for their constructive criticism about the manuscript.

## Author contribution

Conceptualization: ABN EB PNC AA JA. Data curation: EB. Formal analysis: ABN. Investigation: ABN EB. Methodology: ABN EB PNC AA JA. Writing ABN EB.

## Data availability

The complete analysis with embedded code in the form of an interactive html R notebook is available as supplementary data and online at the following address: https://a-bn.github.io/AntibioFoodTM/

