## Supplementary Data for "Antibiotics and food in the American press"

Antibiotics and food in the American press: A text mining study.


Code 

- Show All Code
- Hide All Code
- Download Rmd

### Antibiotics and food in the American press: A text mining study.

######

- Corpus constitution
- Data loading
- Data Processing
  - Information scrapping and cleaning
- Counting and more
  - Publication chronology
  - Counts and TF-IDF
  - Word count evolution through time
- Analyzes
  - GLM
  - Contextual analysis
    - Syntagma curation

*Antibiotics and food in the American press: A text mining study.*  
Antoine Bridier-Nahmias\(\dagger\), Estera Badau\(\dagger\), Pi Nyvall Collen ,Antoine Andremont, Jocelyne Arquembourg

\(\dagger\): These authors contributed equally to this work

### Corpus constitution

The articles have been searched upon the Factiva database, based on key words and expressions used in conjunction. Terms and expressions researched were the following:

```
antibiotic resistance, antimicrobial resistance, 
antibiotic free or antibiotic-free, antibiotics and food, 
antibiotics and farming, antibiotics and resistant, antibiotics and salmonella, 
salmonella and resistant, salmonella and outbreak, 
antibiotics and campylobacter and resistant, antibiotics and routine, 
antibiotics and routinely, antibiotics and One Health;
(antibio* near3 food) or (antibio* near3 farm*) or (antibio* near3 salmonell*) 
or (antibio* near3 campylobacter*) or (antibio* near3 animal*) or 
(antibio* near3 feed)
```

### Data loading

The corpus is consituted by articles saved in independent pdf files.


```
# list pdf files
pdf_list <-
  list.files(path = "../data/corpus/", pattern = "^.*pdf$", full.names = TRUE, recursive = TRUE)
# scrap their text content
pdf_txt <-
  lapply(X = pdf_list, FUN = function(x) paste0(pdf_text(x), collapse = "\n" ))
# split them over each newline
pdf_split <-
  sapply(X = pdf_txt, function(x) str_split(string = unlist(x), pattern = "\n"))
# eliminating leading spaces in each line
pdf_split_strp <-
  lapply(X = pdf_split, function(x) str_replace(string = x, pattern = "^[[:space:]]+", replacement = ""))
```

### Data Processing

#### Information scrapping and cleaning


```
# This function will take care of the parsing
gogo_gadgeto_get_info <- function(text_vector){
  # HD is the title tag
  title <- gsub(pattern = "HD(.*)", replacement = "\\1",
                x = grep(pattern = "^HD.*", x = text_vector, value = TRUE))
  title <- str_remove_all(string = title, pattern = "^ *")
  # BY is the author tag
  author <- gsub(pattern = "(?:BY|By)(.*)", replacement = "\\1",
                 x = grep(pattern = "^(?:BY|By).*", x = text_vector, value = TRUE))
  author <- ifelse(test = length(author) == 0, yes = "", no = author)
  
  # SN is the journal tag
  journal <- gsub(pattern = "SN(.*)", replacement = "\\1",
                  x = grep(pattern = "^SN.*", x = text_vector, value = TRUE))
  journal <- ifelse(test = length(journal) == 0, yes = "", no = journal)
  journal <- str_replace(journal, "^ +", "")
  
  # PD is the publication date tag
  pub_date <- gsub(pattern = "PD(.*)", replacement = "\\1",
                   x = grep(pattern = "^PD.*", x = text_vector, value = TRUE))
  pub_date <- ifelse(test = length(pub_date) == 0, yes = "", no = pub_date)
  pub_date <- gsub(pattern = "aot", replacement = "august", x = pub_date)
  pub_date <- dmy(pub_date)
  
  # output is a tibble with all the informations
  article_info <- tibble(title = title,
                             author = author,
                             journal = journal,
                             pub_date = pub_date)
  return(article_info)
}
# extract informations of each article w/ gogo_gadgeto_get_info
pdf_info <-
  lapply(X = pdf_split_strp, FUN = gogo_gadgeto_get_info)
```


We will now fuse the articles and their respective informations in a dataframe, and then we will remove the headers and footers. This operation is noisy because of the inconsistency in the footer formatting.


```
# making a data.frame with the info and the text
pdf_txt_info <- list()
for (i in 1:length(pdf_split_strp)) {
  pdf_txt_info[[i]] <- 
    cbind.data.frame(pdf_info[[i]], 
                     text = as.character(pdf_split_strp[[i]]), 
                     stringsAsFactors = FALSE)
}
# Removing header and footer in each dataframe
# The footer is inconsistent across article and 
# many different lines are needed to purge it out
# check for a string : sum(unlist(lapply(pdf_txt_info, function(x) str_detect(string = x$text, pattern = "^LP"))))
behead_and_befoot <- function(df_in){
  df_out <-
    df_in %>%
    filter(cumsum(str_detect(text, pattern = "^LP")) >= 1) %>%
    filter(cumsum(str_detect(text, pattern = "^NS") ) < 1) %>%
    filter(cumsum(str_detect(text, pattern = "^Illustrations:") ) < 1) %>%
    filter(cumsum(str_detect(text, pattern = "^ART") ) < 1) %>%
    filter(cumsum(str_detect(text, pattern = "^CT") ) < 1) %>%
    filter(cumsum(str_detect(text, pattern = "^IPD") ) < 1) %>%
    filter(cumsum(str_detect(text, pattern = "^.*\\|.*\\|.*")) < 1) %>%
    filter(cumsum(str_detect(text, pattern = "^AN ")) < 1) %>% 
    filter(cumsum(str_detect(text, pattern = "^RF ")) < 1) %>% 
    filter(cumsum(str_detect(text, pattern = "^CO ")) < 1) %>%
    filter(!str_detect(text, pattern = "Factiva")) %>%
    filter(!str_detect(text, pattern = "^TD$")) %>%
    filter(!str_detect(text, pattern = "^LP$")) %>%
    filter(!str_detect(text, pattern = "^$")) %>% 
    identity()
  return(df_out)
}
txt_clean <-
  lapply(X = pdf_txt_info, 
         FUN = behead_and_befoot)
```


We are ready to unite everything in one data.frame, beforehand we’ll just add a unique id for each article.


```
# before uniting them, each article needs to receive a unique ID
for (i in 1:length(txt_clean)) {
  txt_clean[[i]] <- cbind.data.frame(txt_clean[[i]], id = i)
  txt_clean[[i]]$text[1] <- paste(txt_clean[[i]]$title[1], 
                                  txt_clean[[i]]$text[1], 
                                  sep = " ")
}
# uniting everything into a big dataframe
corpus_txt <-
  do.call(rbind, txt_clean)
```


The tokenzation can now take place. We can use multiple ngrams size, we will start with 1grams first *i.e*: words.


```
corpus_1_grams_unfiltered <- 
  unnest_tokens(tbl = corpus_txt, 
                output = "word", 
                input = "text", 
                token = "ngrams", 
                n = 1)
```

### Counting and more

Let’s first extract some figures about the whole corpus


```
articla <-
  length(unique(corpus_1_grams_unfiltered$id))
articla_by_journal <-
  corpus_1_grams_unfiltered %>%
  group_by(journal) %>%
  summarize(n = n_distinct(id))
worda <- 
  format(nrow(corpus_1_grams_unfiltered), big.mark = ",")
worda_uniq <-
  format(length(unique(corpus_1_grams_unfiltered$word)), big.mark = ",")
worda_by_journal <-
  corpus_1_grams_unfiltered %>%
  group_by(journal) %>% count()
worda_uniq_by_journal <-
  corpus_1_grams_unfiltered %>%
  group_by(journal) %>%
  summarize(n = n_distinct(word))
# articla;articla_by_journal;worda;worda_uniq; worda_by_journal;worda_uniq_by_journal
```


- The total number of articles:
- overall: 516
- by journal:
- The total number of words in the corpus
- overall: 493,202
- by journal:
- The total number of distinct words in the corpus
- overall: 22,814
- by journal:

#### Publication chronology


```
timeline_events <- read_delim("../data/timeline_events.tsv", 
                              "\t", escape_double = FALSE, trim_ws = TRUE, comment = "#")
```


```
Parsed with column specification:
cols(
  date = [34mcol_date(format = "")[39m,
  full_event = [31mcol_character()[39m,
  event_label = [31mcol_character()[39m
)
```


```
timeline_events <-
  timeline_events %>%
  mutate(event_label = str_replace_all(string = event_label, pattern = "XXX", replacement = "\n")) %>%
  mutate(date = ymd(date)) %>%
  mutate(date_lab = paste(month(date, label = TRUE, abbr = TRUE, locale = "en_US.utf8"), year(date))) %>%
  mutate(event_label = paste0(event_label,"\n", date_lab)) %>% 
  mutate(ypos = 0.1, ypos = ypos * c(1, -1)) %>%  # in order to appear above or under the timeline
  mutate(ydate = ypos*0.5)
```


```
longer object length is not a multiple of shorter object length
```


```
article_hist_dodge <-
  corpus_1_grams_unfiltered %>%
  ungroup() %>%
  select(id, pub_date, journal) %>%
  group_by(id) %>%
  slice(1) %>%
  mutate(year = ymd(paste0(year(pub_date),"01","01"))) %>%
  group_by(year, journal) %>%
  mutate(count = n()) %>%
  slice(1)
histo <-
  ggplot(data = article_hist_dodge) +
  geom_col(mapping = aes(x = year, y = count, fill = journal), 
           position = "dodge", alpha = 0.8, colour = "black") +
  scale_color_manual(values = c(alpha("black", 0.1), alpha("black", 0.1))) +
  scale_fill_manual(values = c("steelblue", "violetred3")) +
  scale_x_date(breaks = seq(from = ymd("1980/01/01"), to = ymd("2015/01/01"), by = "5 years"),
               date_labels = "%Y",
               minor_breaks = waiver(),
               date_minor_breaks = "1 years",
               limits = c(ymd("1979/01/01"),ymd("2017/01/01"))) + 
  background_grid(major = "xy", 
                  minor = "xy", 
                  colour.major = rgb(red = 0.5,green = 0.5,blue = 0.5, alpha = 0.5),
                  colour.minor = rgb(red = 0.5,green = 0.5,blue = 0.5, alpha = 0.1)) +
  theme(legend.position = "top",
        legend.direction = "horizontal")
histo
```


```
timeline <- 
  ggplot(data = timeline_events) +
  geom_point(mapping = aes(x = date, y = ypos)) +
  geom_segment(mapping = aes(xend = date, x = date, y = ypos, yend = 0)) +
  geom_hline(yintercept = 0, color = "black", size=0.3) + # timeline itself
  ggrepel::geom_label_repel(mapping = aes(x = date, y = ypos, label = event_label), inherit.aes = FALSE) +
  # ggrepel::geom_label_repel(mapping = aes(x = date, y = ydate, label = date_lab, angle = 25), point.padding = 0, inherit.aes = FALSE) +
  scale_x_date(breaks = seq(from = ymd("1980/01/01"), to = ymd("2015/01/01"), by = "5 years"),
               date_labels = "%Y",
               minor_breaks = waiver(),
               date_minor_breaks = "1 years",
               limits = c(ymd("1979/01/01"),ymd("2017/01/01"))) + 
  theme(axis.line.y = element_blank(),
        axis.text.y = element_blank(),
        axis.title.x = element_blank(),
        axis.title.y = element_blank(),
        axis.ticks.y = element_blank(),
        axis.text.x = element_blank(),
        axis.ticks.x = element_blank(),
        axis.line.x = element_blank()
          )
timeline
```

#### Counts and TF-IDF

We will compute the following:

- *corpus\_length*: the total number of documents in the corpus
- *article\_length*: the total number of words in each article
- *n\_article*: the count of each word in each article
- *word\_in\_n*: for each word, how many articles contain it
- *n\_tot*: total count of each word
- *n\_journal*: total count of each word by journal


```
suppressMessages(
  my_stop_words <-
    read_table(
      file = "../data/my_stop_words.txt",
      col_names = "stop_word"
    )
)
corpus_tfidf_full <-
  corpus_1_grams_unfiltered %>%
  # filter(id %in% c(30:100,400:500) ) %>%
  # filter(id %in% c(517) ) %>%
  
  # clean dataset
  mutate(word = str_to_lower(string = word)) %>% 
  mutate(word = str_replace_all(string = word, pattern = "([[:alpha:]]*)\\.([[:alpha:]]*)", replacement = "\\1\\2")) %>%
  mutate(word = str_replace_all(string = word, pattern = "'s$", replacement = "")) %>%
  # mutate(word = str_replace_all(string = word, pattern = "[^a-z]", replacement = "")) %>%
  filter(!(str_detect(string = word, pattern = "washpostcom"))) %>% # Tokenization splits on @ !!!!!!!
  filter(word != "") %>% 
  filter(nchar(word) > 1) %>% 
  filter(!str_detect(string = word, pattern = "^[0-9]|[[:punct:]]+$")) %>%
  
  # lemmatization (better than stemming) and last filtering
  mutate(stem = stem_words(word)) %>%
  filter(!word %in% my_stop_words$stop_word) %>% 
  
  # total number of articles
  mutate(corpus_length = length(unique(id) )) %>%  
  
  # total word in article
  group_by(id) %>%
  mutate(article_length = n()) %>%
  
  # count of each word by article
  group_by(id, stem) %>% 
  mutate(n_article = n()) %>% 
  
  # word count
  group_by(stem) %>% 
  mutate(n_total = n()) %>% 
  group_by(stem, journal) %>% 
  mutate(n_journal = n()) %>% 
  ungroup() %>%
  # compute tf-idf
  
  mutate(tf = n_article / article_length) %>% # text frequency
  group_by(stem) %>% 
  mutate(word_in_n = length(unique(id))) %>% 
  mutate(idf = log(corpus_length / word_in_n) ) %>% # inverse document frequency
  mutate(tf_idf = tf * idf) %>% 
  ungroup() %>% 
  
  # Choose a representant for each stem, the most common term could be the best
  group_by(stem) %>%
  mutate(ori_word = word) %>%
  group_by(ori_word) %>% 
  mutate(n_ori = n()) %>%
  arrange(desc(n_ori)) %>%
  group_by(stem) %>% 
  mutate(word = word[1]) %>%
  select(-n_ori) %>% 
  ungroup()
# write_delim(x = corpus_tfidf_full, path = "../output/corpus_tfidf_full.tsv", delim = "\t", col_names = TRUE)
# corpus_tfidf_full <- 
#   read_delim(file = "../output/corpus_tfidf_full.tsv", delim = "\t", col_names = TRUE)
corpus_tfidf <-
  corpus_tfidf_full %>% 
  # reduce
  group_by(id, word) %>%
  slice(1) %>% 
  ungroup() %>%
  
  # filter out  words with tf-idf == 0
  filter(tf_idf > 0) %>% 
  identity()
corpus_tfidf %>% 
  filter(!(word %in% stop_words$word)) %>%
  filter(n_total >= 50) %>% 
  arrange(desc(n_total)) %>% 
  group_by(word, journal) %>% 
  slice(1) %>%
  select(word, journal, pub_date, n_total, n_journal) %>%
  arrange(desc(n_total)) %>% 
  datatable(caption = "Words appearing at least 50 times", filter = "top") %>% 
  identity()
```

#### Word count evolution through time


```
corpus_year <-
  corpus_tfidf_full %>%
  # Filter out a word if one member of the family (ori_word) is a stopword
  group_by(word) %>% 
  mutate(is_stop = ifelse(test = sum(ori_word %in% stop_words$word) >= 1, yes = TRUE, no = FALSE)) %>%
  filter(!is_stop) %>% 
  mutate(is_stop = NULL) %>% 
  ungroup() %>% 
  filter(n_total > 350) %>%
  mutate(year = year(pub_date)) %>%
  group_by(word, year, journal) %>%
  mutate(n_year = n()) %>%
  slice(1) %>%
  ungroup() %>%
  mutate(word = reorder(word, desc(n_total)))# Conversion to factor for ordering in the facet_wrapping of the plot
my_labeller <-
  unique(paste0(corpus_year$word, "(",corpus_year$n_total,")"))
names(my_labeller) <- unique(corpus_year$word)
word_time_plot <-
  ggplot(corpus_year) +
  geom_line(mapping = aes(x = pub_date, y = n_year, colour = journal)) +
  ylab("count") +
    xlab("Year") +
  scale_color_manual(values = c("steelblue", "violetred3")) +
  scale_x_date(breaks = seq(from = ymd("1980/01/01"), to = ymd("2015/01/01"), by = "5 years"),
               date_labels = "%Y",
               minor_breaks = waiver(),
               date_minor_breaks = "1 years",
               limits = c(ymd("1979/01/01"),ymd("2017/01/01"))) + # 1 year before because dodge needs a bit of space apparently
  theme_bw() +
  theme(legend.position = "top", legend.text = element_text(size = 12)) +
  facet_wrap(~word, ncol = 3, labeller = as_labeller(my_labeller), scales = "free")
# ggdraw() + draw_plot(word_time_plot) + ggsave(filename = "../output/words_time.pdf", width = 30, height = 90, units = "cm")
word_time_plot
```


```
# For the paper
selected_word_time_plot <-
  corpus_year %>%
  filter(str_detect(string = word, pattern = "^antibiotics$|^farm$|^industry$")) %>%
  ggplot() +
  geom_line(mapping = aes(x = pub_date, y = n_year, colour = journal)) +
  ylab("count") +
  xlab("Year") +
  scale_color_manual(values = c("steelblue", "violetred3"), 
                     guide = guide_legend(title = NULL)) +
  scale_x_date(breaks = seq(from = ymd("1980/01/01"), to = ymd("2015/01/01"), by = "5 years"),
               date_labels = "%Y",
               minor_breaks = waiver(),
               date_minor_breaks = "1 years",
               limits = c(ymd("1979/01/01"),ymd("2017/01/01"))) + # 1 year before because dodge needs a bit of space apparently
  theme_bw() +
  theme(legend.position = "top", legend.text = element_text(size = 12)) +
  facet_wrap(~word, ncol = 1, labeller = as_labeller(my_labeller), scales = "free")
```


```
selected_word_time_plot
```

### Analyzes

#### GLM

What are the term that could discriminate between an article from the WP and the NYT?


```
corpus_glm_wide <-
  corpus_tfidf %>% 
  select(word, id, journal, tf_idf) %>% 
  mutate(id_journal = journal, journal = NULL) %>% # journal is a word present in the corpus 
  spread(key = word, value = tf_idf) %>% 
  base::replace(x = ., list = is.na(.), values = 0) %>%
  ungroup() %>% 
  select(-id) %>% 
  mutate(id_journal = as.factor(id_journal)) %>% 
  ungroup() %>% 
  identity()
standardize_classic <- 
  function(x) return((x - mean(x)) / sd(x))
standardize_gelman <- 
  function(x) return(x - mean(x) / 2 * sd(x)) # strange, see https://andrewgelman.com/2009/07/11/when_to_standar/
standardized_tf_idf <- # corpus_glm_wide[, -1]
  apply(X = as.matrix(corpus_glm_wide[ ,-1]), 
        MARGIN = 2,  
         FUN = standardize_classic)
response_var <- 
  corpus_glm_wide$id_journal
response_var_bool <- 
  ifelse(test = response_var == "The Washington Post", 
         yes = TRUE, 
         no = FALSE)
```


```
set.seed(c(12,10,2018,18,20)) # cv.glmnet has a random part
system.time(
corpus_lasso <- 
  cv.glmnet(x = standardized_tf_idf, 
                          y = response_var_bool, 
                          family = "binomial", 
                          nfolds = 10, 
                          type.measure = "auc", # could be "auc" or "class"
                          intercept = FALSE,
                          alpha = 0.5)
)
```


```
   user  system elapsed 
 13.482   0.293  13.802
```


```
plot(corpus_lasso)
```


```
worda <- 
  dimnames(coef(corpus_lasso))[[1]]
beta_df <- 
  data.frame(word = worda,beta_coef = as.vector(coef.cv.glmnet(corpus_lasso, corpus_lasso$lambda.min))) %>% 
  filter(beta_coef != 0) %>%
  arrange(desc(beta_coef)) %>% 
  mutate(p = exp((beta_coef)) / (1 + exp((beta_coef))) ) %>% 
  mutate(lab_beta_coef = (signif(beta_coef, 2))) %>% 
  identity()
beta_df
```


```
tilos <-
  ggplot(data = beta_df, mapping = aes(x = factor(0), y = reorder(word, beta_coef))) +
  geom_tile(aes(fill = beta_coef)) +
  geom_text(aes(label = word)) +
  scale_fill_gradient2(low = "steelblue",
                       mid = "white", 
                       high = "violetred3", 
                       midpoint = 0, 
                       space = "Lab", 
                       breaks = c(max(beta_df$beta_coef), min(beta_df$beta_coef)),
                       labels = c("WP", "NYT") ) +
  scale_y_discrete(labels = (sort(beta_df$lab_beta_coef, decreasing = FALSE)) ) +
  ggtitle("") +
  xlab("word") +
  ylab("beta coefficient from the standardized logistic regression") +
  theme_classic() +
  theme(axis.text = element_text(), 
        axis.text.x = element_blank(), 
        axis.ticks.x = element_blank(), 
        panel.background = element_blank(), 
        panel.grid.major = element_blank(), 
        panel.grid.minor = element_blank(),
        legend.title = element_blank()
        ) +
  geom_blank()
tilos
```


```
# predos <- 
#   predict.cv.glmnet(object = corpus_lasso, 
#                     newx = standardized_tf_idf, 
#                     s = "lambda.min", 
#                     type = "class")
# table(predos == response_var_bool)
```

#### Contextual analysis

##### Syntagma curation

In the next section, we will concentrate on counting manually curated terms or expressions (syntagmas) They will be presented in a named list containing all the terms considered equivalent. We will then extract all the sentences in which they occur and analyse their context (in general and across time).


```
curated_syntagma <- 
  list(
    "antibiotic_resistance" = 
      c("antibiotic resistance", "antibiotic-resistance", "resistant to antibiotics", "resistance to antibiotics"),
    "antibiotic_free" = 
      c("antibiotic free", "antibiotic-free", "antibioticsfree", "free of antibiotics"),
    "routine_use" = 
      c("routine use", "routinely used"),
    "judicious_use" = 
      c("judicious use"),
    "responsible_use" = 
      c("responsible use"),
    "prudent_use" = 
      c("prudent use"),
    "indiscriminate_use" = 
      c("indiscriminate use"),
    "food_borne" = 
      c("food borne", "food-borne")
  )
curated_syntagma
```


```
$antibiotic_resistance
[1] "antibiotic resistance"     "antibiotic-resistance"     "resistant to antibiotics"  "resistance to antibiotics"

$antibiotic_free
[1] "antibiotic free"     "antibiotic-free"     "antibioticsfree"     "free of antibiotics"

$routine_use
[1] "routine use"    "routinely used"

$judicious_use
[1] "judicious use"

$responsible_use
[1] "responsible use"

$prudent_use
[1] "prudent use"

$indiscriminate_use
[1] "indiscriminate use"

$food_borne
[1] "food borne" "food-borne"
```


The first step is to divide the corpus in sentences.  
Remark `unnest_tokens(token = "sentences")` clearly fails whenever it encounters an abbreviation containing a dot.


```
corpus_sentences <-
  corpus_txt %>% 
  # Each article has to be re-concatenated
  group_by(id) %>% 
  # filter(id %in% 1:5) %>%
  mutate(article = paste(text, collapse = " ")) %>%
  select(-text) %>% 
  distinct() %>%
  ungroup() %>% 
# Could/should be done in one pass with a list of terms!
  mutate(article = str_replace_all(string = article, pattern = regex(pattern = "dr\\.", ignore_case = TRUE), replacement = "dr")) %>%
  mutate(article = str_replace_all(string = article, pattern = regex(pattern = "prof\\.", ignore_case = TRUE), replacement = "prof")) %>%
  mutate(article = str_replace_all(string = article, pattern = regex(pattern = "mr\\.", ignore_case = TRUE), replacement = "mr")) %>%
  mutate(article = str_replace_all(string = article, pattern = regex(pattern = "ms\\.", ignore_case = TRUE), replacement = "ms")) %>%
  mutate(article = str_replace_all(string = article, pattern = regex(pattern = "mrs\\.", ignore_case = TRUE), replacement = "mrs")) %>%
  mutate(article = str_replace_all(string = article, pattern = regex(pattern = "st\\.", ignore_case = TRUE), replacement = "st")) %>%
  mutate(article = str_replace_all(string = article, pattern = regex(pattern = "rep\\.", ignore_case = TRUE), replacement = "rep")) %>%
  mutate(article = str_replace_all(string = article, pattern = regex(pattern = "u\\.s\\.", ignore_case = TRUE), replacement = "usa")) %>%
  mutate(article = str_replace_all(string = article, pattern = regex(pattern = "f\\.d\\.a\\.", ignore_case = TRUE), replacement = "fda")) %>%
  mutate(article = str_replace_all(string = article, pattern = regex(pattern = "gov\\.", ignore_case = TRUE), replacement = "gov")) %>%
  mutate(article = str_replace_all(string = article, pattern = regex(pattern = "sen\\.", ignore_case = TRUE), replacement = "sen")) %>%
  mutate(article = str_replace_all(string = article, pattern = regex(pattern = "( .{1})\\.", ignore_case = TRUE), replacement = "\\1")) %>%
  unnest_tokens(output = "sentence", 
                input = article, 
                token = "sentences", 
                to_lower = TRUE) %>%
  mutate(length = nchar(sentence)) %>%
  select(length, everything()) %>% 
  ungroup() %>% 
  identity()
# Diagnose problems with abbreviations (Mr. Dr. etc)
end_sentences_corpus <-
  corpus_sentences %>%
  filter(str_detect(string = sentence, pattern = "^[[:alnum:]]{1,5}\\.$")) %>%
  group_by(sentence) %>%
  summarize(n = n()) %>%
  ungroup() %>%
  mutate(n_char = nchar(sentence)) %>%
  arrange(desc(n))
```


We can now try to isolate sentences containing our terms of interest.


```
corpus_syntagma_sentence <-
  lapply(X = curated_syntagma, FUN = function(syntagma){
    corpus_sentences %>%
      rowwise() %>%
      mutate(syntagmus = ifelse(test = sum(str_detect(string = sentence, pattern = syntagma)) > 0, yes = syntagma, no = NA)) %>%
      filter(!is.na(syntagmus))
  })
corpus_syntagma_sentence <-
  do.call(what = rbind, args = corpus_syntagma_sentence)
table(corpus_syntagma_sentence$syntagmus)
```


```
      antibiotic free antibiotic resistance            food borne    indiscriminate use         judicious use           prudent use 
                  125                   299                   119                    16                    14                     7 
      responsible use           routine use 
                    1                    53
```


```
table(corpus_syntagma_sentence$syntagmus, corpus_syntagma_sentence$journal)
```


```
                        The New York Times The Washington Post
  antibiotic free                       88                  37
  antibiotic resistance                165                 134
  food borne                            55                  64
  indiscriminate use                    13                   3
  judicious use                          9                   5
  prudent use                            6                   1
  responsible use                        1                   0
  routine use                           38                  15
```


On this new dataframe, we can count the occurence of each word in the sentence context of each syntagma, overall and divided by journal:


```
corpus_syntagma_word <-
  corpus_syntagma_sentence %>%
  unnest_tokens(output = "word", input = sentence, token = "ngrams", n = 1) %>%
  filter(!(word %in% stop_words$word)) %>%
  filter(is.na(str_match(string = word, pattern = "[0-9]"))) %>% # no match in str_match() returns NA
  mutate(word = str_replace_all(string = word, pattern = "\\.", replacement = "")) %>%
  mutate(word = str_replace_all(string = word, pattern = "(.*)'s", replacement = "\\1")) %>%
  mutate(stem = stem_words(word)) %>%
  
  # Choose a representant for each stem, the most common term could be the best
  group_by(stem) %>%
  mutate(ori_word = word) %>%
  group_by(ori_word) %>% 
  mutate(n_ori = n()) %>%
  arrange(desc(n_ori)) %>%
  group_by(stem) %>% 
  mutate(word = word[1]) %>%
  select(-n_ori) %>% 
  ungroup() %>% 
  
  group_by(syntagmus, stem) %>%
  mutate(n_total = n()) %>%
  group_by(syntagmus, journal, stem) %>%
  mutate(n_journal = n()) %>%
  slice(1) %>%
  ungroup() %>% 
  select(syntagmus, word, journal, n_journal, n_total) %>%
  arrange(desc(n_total)) %>%
  identity()
```


```
Grouping rowwise data frame strips rowwise natureGrouping rowwise data frame strips rowwise nature
```


```
corpus_syntagma_word %>% 
    datatable(filter = "top", rownames = FALSE, options = list(pageLength = 10))
```


Figures printing and saving


Packages loading


```
─ Session info ─────────────────────────────────────────────────────────────────────────────────────────────────────────────────────────────

─ Packages ─────────────────────────────────────────────────────────────────────────────────────────────────────────────────────────────────
 package        * version date       lib source        
 askpass          1.1     2019-01-13 [1] CRAN (R 3.6.0)
 assertthat       0.2.1   2019-03-21 [1] CRAN (R 3.6.0)
 backports        1.1.4   2019-04-10 [1] CRAN (R 3.6.0)
 base64enc        0.1-3   2015-07-28 [1] CRAN (R 3.6.0)
 broom            0.5.2   2019-04-07 [1] CRAN (R 3.6.0)
 callr            3.2.0   2019-03-15 [1] CRAN (R 3.6.0)
 cli              1.1.0   2019-03-19 [1] CRAN (R 3.6.0)
 codetools        0.2-16  2018-12-24 [2] CRAN (R 3.6.0)
 colorspace       1.4-1   2019-03-18 [1] CRAN (R 3.6.0)
 cowplot        * 0.9.4   2019-01-08 [1] CRAN (R 3.6.0)
 crayon           1.3.4   2017-09-16 [1] CRAN (R 3.6.0)
 crosstalk        1.0.0   2016-12-21 [1] CRAN (R 3.6.0)
 data.table       1.12.2  2019-04-07 [1] CRAN (R 3.6.0)
 desc             1.2.0   2018-05-01 [1] CRAN (R 3.6.0)
 devtools         2.0.2   2019-04-08 [1] CRAN (R 3.6.0)
 digest           0.6.18  2018-10-10 [1] CRAN (R 3.6.0)
 dplyr          * 0.8.0.1 2019-02-15 [1] CRAN (R 3.6.0)
 DT             * 0.5     2018-11-05 [1] CRAN (R 3.6.0)
 evaluate         0.13    2019-02-12 [1] CRAN (R 3.6.0)
 foreach        * 1.4.4   2017-12-12 [1] CRAN (R 3.6.0)
 fs               1.2.7   2019-03-19 [1] CRAN (R 3.6.0)
 generics         0.0.2   2018-11-29 [1] CRAN (R 3.6.0)
 ggplot2        * 3.1.1   2019-04-07 [1] CRAN (R 3.6.0)
 ggrepel          0.8.0   2018-05-09 [1] CRAN (R 3.6.0)
 glmnet         * 2.0-16  2018-04-02 [1] CRAN (R 3.6.0)
 glue             1.3.1   2019-03-12 [1] CRAN (R 3.6.0)
 gtable           0.3.0   2019-03-25 [1] CRAN (R 3.6.0)
 hms              0.4.2   2018-03-10 [1] CRAN (R 3.6.0)
 htmltools        0.3.6   2017-04-28 [1] CRAN (R 3.6.0)
 htmlwidgets      1.3     2018-09-30 [1] CRAN (R 3.6.0)
 httpuv           1.5.1   2019-04-05 [1] CRAN (R 3.6.0)
 iterators        1.0.10  2018-07-13 [1] CRAN (R 3.6.0)
 janeaustenr      0.1.5   2017-06-10 [1] CRAN (R 3.6.0)
 jsonlite         1.6     2018-12-07 [1] CRAN (R 3.6.0)
 knitr            1.22    2019-03-08 [1] CRAN (R 3.6.0)
 koRpus         * 0.11-5  2018-10-28 [1] CRAN (R 3.6.0)
 koRpus.lang.en * 0.1-2   2018-03-21 [1] CRAN (R 3.6.0)
 labeling         0.3     2014-08-23 [1] CRAN (R 3.6.0)
 later            0.8.0   2019-02-11 [1] CRAN (R 3.6.0)
 lattice          0.20-38 2018-11-04 [2] CRAN (R 3.6.0)
 lazyeval         0.2.2   2019-03-15 [1] CRAN (R 3.6.0)
 lubridate      * 1.7.4   2018-04-11 [1] CRAN (R 3.6.0)
 magrittr         1.5     2014-11-22 [1] CRAN (R 3.6.0)
 Matrix         * 1.2-17  2019-03-22 [2] CRAN (R 3.6.0)
 memoise          1.1.0   2017-04-21 [1] CRAN (R 3.6.0)
 mime             0.6     2018-10-05 [1] CRAN (R 3.6.0)
 munsell          0.5.0   2018-06-12 [1] CRAN (R 3.6.0)
 nlme             3.1-139 2019-04-09 [2] CRAN (R 3.6.0)
 pdftools       * 2.2     2019-03-10 [1] CRAN (R 3.6.0)
 pillar           1.3.1   2018-12-15 [1] CRAN (R 3.6.0)
 pkgbuild         1.0.3   2019-03-20 [1] CRAN (R 3.6.0)
 pkgconfig        2.0.2   2018-08-16 [1] CRAN (R 3.6.0)
 pkgload          1.0.2   2018-10-29 [1] CRAN (R 3.6.0)
 plyr             1.8.4   2016-06-08 [1] CRAN (R 3.6.0)
 prettyunits      1.0.2   2015-07-13 [1] CRAN (R 3.6.0)
 processx         3.3.0   2019-03-10 [1] CRAN (R 3.6.0)
 promises         1.0.1   2018-04-13 [1] CRAN (R 3.6.0)
 ps               1.3.0   2018-12-21 [1] CRAN (R 3.6.0)
 purrr            0.3.2   2019-03-15 [1] CRAN (R 3.6.0)
 qpdf             1.1     2019-03-07 [1] CRAN (R 3.6.0)
 R6               2.4.0   2019-02-14 [1] CRAN (R 3.6.0)
 Rcpp             1.0.1   2019-03-17 [1] CRAN (R 3.6.0)
 readr          * 1.3.1   2018-12-21 [1] CRAN (R 3.6.0)
 remotes          2.0.4   2019-04-10 [1] CRAN (R 3.6.0)
 rlang            0.3.4   2019-04-07 [1] CRAN (R 3.6.0)
 rmarkdown        1.12    2019-03-14 [1] CRAN (R 3.6.0)
 rprojroot        1.3-2   2018-01-03 [1] CRAN (R 3.6.0)
 rstudioapi       0.10    2019-03-19 [1] CRAN (R 3.6.0)
 scales           1.0.0   2018-08-09 [1] CRAN (R 3.6.0)
 sessioninfo      1.1.1   2018-11-05 [1] CRAN (R 3.6.0)
 shiny            1.3.2   2019-04-22 [1] CRAN (R 3.6.0)
 SnowballC        0.6.0   2019-01-15 [1] CRAN (R 3.6.0)
 stringi          1.4.3   2019-03-12 [1] CRAN (R 3.6.0)
 stringr        * 1.4.0   2019-02-10 [1] CRAN (R 3.6.0)
 sylly          * 0.1-5   2018-07-29 [1] CRAN (R 3.6.0)
 sylly.en         0.1-3   2018-03-19 [1] CRAN (R 3.6.0)
 textstem       * 0.1.4   2018-04-09 [1] CRAN (R 3.6.0)
 tibble           2.1.1   2019-03-16 [1] CRAN (R 3.6.0)
 tidyr          * 0.8.3   2019-03-01 [1] CRAN (R 3.6.0)
 tidyselect       0.2.5   2018-10-11 [1] CRAN (R 3.6.0)
 tidytext       * 0.2.0   2018-10-17 [1] CRAN (R 3.6.0)
 tokenizers       0.2.1   2018-03-29 [1] CRAN (R 3.6.0)
 usethis          1.5.0   2019-04-07 [1] CRAN (R 3.6.0)
 withr            2.1.2   2018-03-15 [1] CRAN (R 3.6.0)
 xfun             0.6     2019-04-02 [1] CRAN (R 3.6.0)
 xtable           1.8-4   2019-04-21 [1] CRAN (R 3.6.0)
 yaml             2.2.0   2018-07-25 [1] CRAN (R 3.6.0)

[1] /home/abn/R/x86_64-pc-linux-gnu-library/3.6
[2] /usr/lib/R/library
```
